# Immune evasion and immunotherapy resistance via TGF-beta activation of extracellular matrix genes in cancer associated fibroblasts

**DOI:** 10.1101/265884

**Authors:** Ankur Chakravarthy, Lubaba Khan, Nathan Peter Bensler, Pinaki Bose, Daniel D. De Carvalho

## Abstract

**The ability to disseminate, invade and successfully colonise other tissues is a critical hallmark of cancer that involves remodelling of the extracellular matrix (ECM) laid down by fibroblasts ^1^. Moreover, Cancer-Associated-Fibroblasts (CAFs) produce key growth factors and cytokines as components of the ECM that fuel tumour growth, metastasis and chemoresistance, and immune response ^2-4^. ECM changes also predict prognosis in pancreatic 5 and colorectal cancers **^6,7^**. Here, we examine the landscape of ECM-gene dysregulation pan-cancer and find that a subset of ECM genes is (*i*) dysregulated specifically in cancer, (*ii*) adversely prognostic, (*iii*) linked to TGF-beta signalling and transcription in Cancer-Associated-Fibroblasts, (*iv*) enriched in immunologically active cancers, and (*v*) predicts responses to Immune checkpoint blockade better than mutation burden, cytolytic activity, or an interferon signature, thus identifying a novel mechanism of immune evasion for patient stratification in precision immunotherapy and pharmacological modulation.**

---

Initially, to study ECM gene dysregulation across cancers, we defined a transcriptional signature to distinguish malignant (n = 8043) and normal samples (n = 704) accounting for tumour type (n = 15) from TCGA and tested for enrichment of an ECM-associated gene-set we curated based on gene ontology terms(Table S1, Figure S1A). This identified 58/239 ECM genes to be cancer-associated (hereby Cancer-associated-ECM genes/C-ECM genes) (Table S2), representing significant enrichment amongst both upregulated (OR = 3.51, p < 3.9e-8) and downregulated (OR = 2.57, p = 3e-5, Fisher’s Exact Test) genes in malignant tissues (Figure 1A). Upon summarisation using ssGSEA (single sample Gene Set Enrichment Analysis) scores^8,9^, these show broad variation across tumour types (Figure 1B, Figure S1B-C). We then performed a Cox regression based on quartile-thresholded C-ECM scores with AJCC stage and tumour-type as strata highlighted to examine the prognostic impact of this dysregulation, which showed upregulated C-ECM genes to be significantly prognostic (Figure 1C-D, HR = 1.73, p < 6.3e-7 for top vs bottom quartile) while downregulated genes were not (Figure S1D), suggesting that the variation we observed in C-ECM gene transcription is clinically relevant. Given the previously identified role of distinct stromal cells in determining the composition and behaviour of the ECM ^10^, we then attempted to infer the potential cell types driving C-ECM transcriptional variation to examine if changes in cellular composition, along with cell-type specific transcriptional changes, could drive C-ECM gene dysregulation using a range of computational approaches, and found multiple indicators that C-ECM gene dysregulation originated in Cancer Associated Fibroblasts.

**Figure 1:**
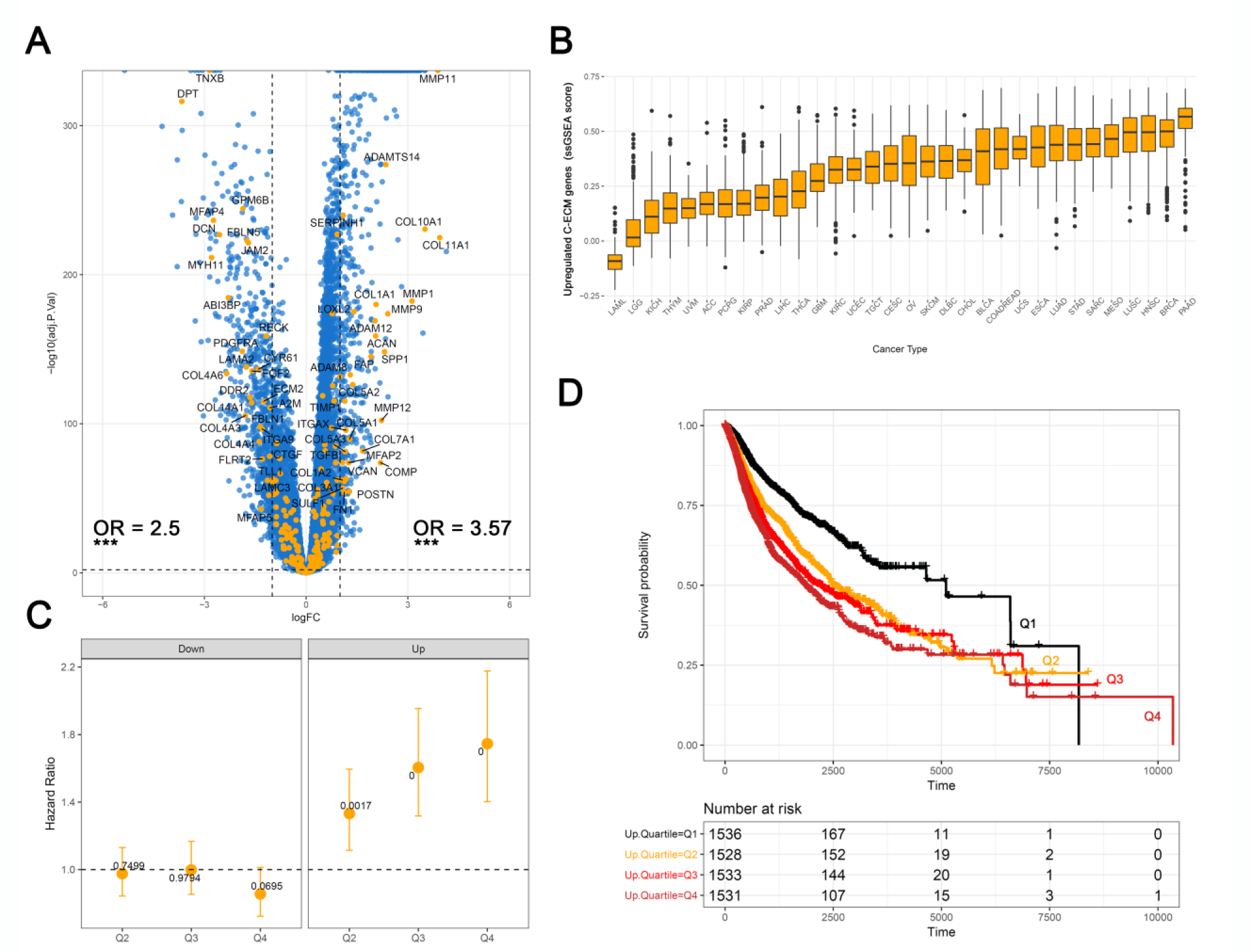
ECM genes are significantly associated with tumourigenesis and are prognostic. **A.** Volcano-plot showing fold changes for genes differentially expressed between cancer and normal. showing ECM gene enrichment for upregulated and downregulated genes. **B.** Boxplots of C-ECM-up enrichment scores show variation across tumour types (S1C for downregulated genes). **C.** Plot of Cox model coefficients by quartile for C-ECM-up and down scores pan-cancer. **D.** Unadjusted Kaplan-Meier curves showing survival by C-ECM-up-quartile.

First, tumour purity estimated using ABSOLUTE ^11^ were inversely correlated for both C-ECM up and down scores (Figure 2A, S2A). Second, projecting the expression signature onto microdissected Ovarian cancer stroma, matched epithelium, and their normal counterparts ^12^ (GSE40595) resulted in clustering by sample type with strong stromal expression (Figure 2B). Additionally, probes differentially expressed between cancer epithelium and stroma, and between cancer and normal stroma, were significantly enriched for both C-ECM-up and down genes (Figure 2C) for the former, and C-ECM-up genes for the latter. Third, deconvolution analysis using MethylCIBERSORT implicated CAFs, CD8 T-cells, and CD14-monocytes as directly correlated with C-ECM signature scores (Figure 2D). Importantly, upregulated C-ECM genes (ssGSEA scores) showed a positive correlation to the inferred CAF frequency in most TCGA cancer types (Figure S2B). We also validated these inferences of cellular association using transcript levels of well-known marker genes (Cytolytic activity (geometric mean of *GZMA, PRF1*) and *CD8A* expression for CD8 T-cells, *ACTA2* for CAFs and *CD14* for monocytes, Figure S2C), whereupon we noticed strong, consistent, agreement.

**Figure 2:**
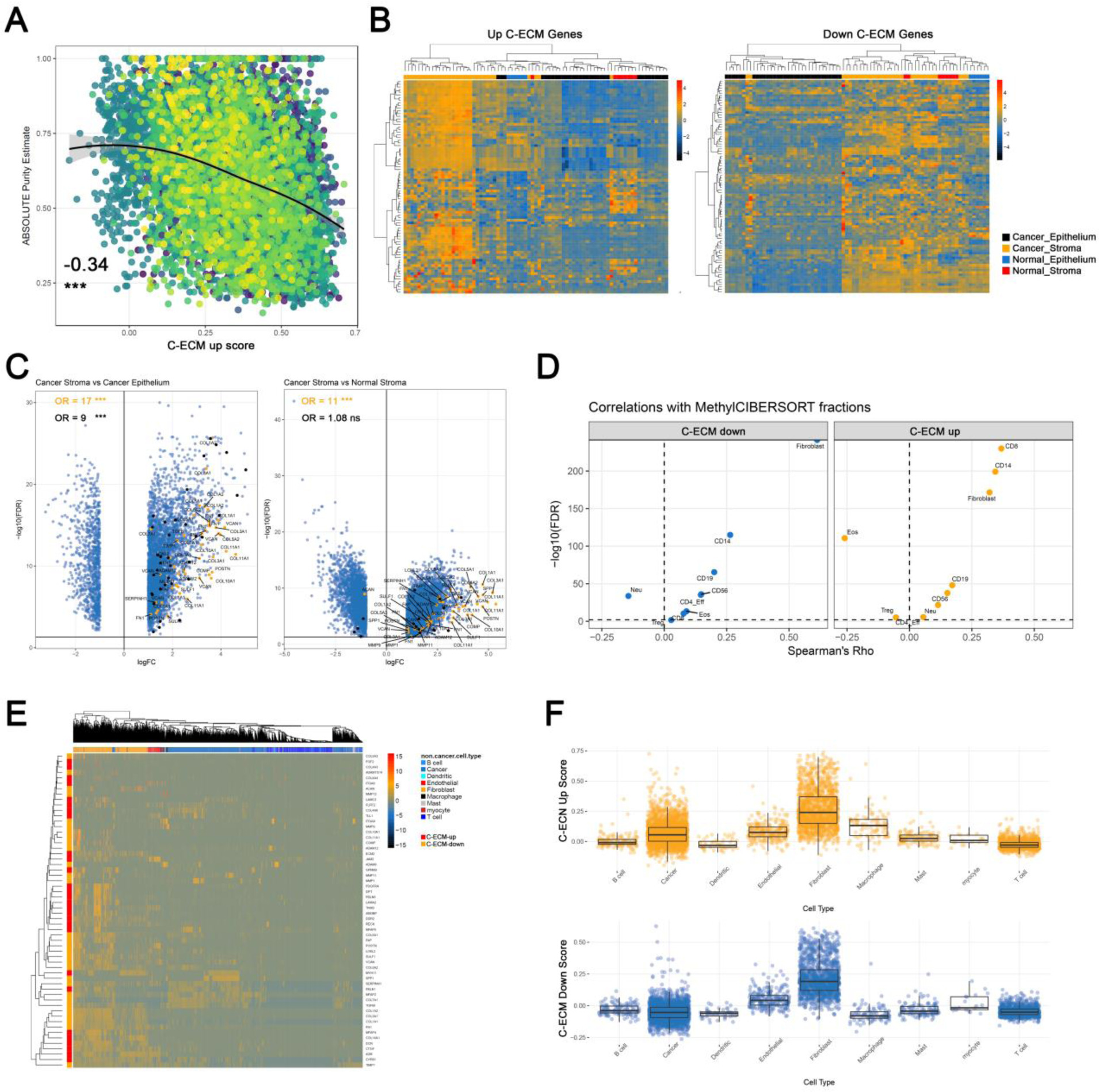
C-ECM transcription is associated with stroma, especially CAFs. **A.** ABSOLUTE purity estimates are inversely correlated with C-ECM-up score, suggesting stromal origin, colours represent cancer types, number shows Spearman’s Rho. **B.** Heatmaps of C-ECM-up and down signatures projected onto epithelium and stroma from ovarian cancers. Rows show expression z-scores, samples are in columns. Annotation bars indicate tissue type. **C.** Volcano-plots show C-ECM genes (upregulated in orange, downregulated in black) in the context of differential expression between cancer stroma and epithelium, and cancer and normal stroma. **D.** Volcano-plots showing Spearman’s correlations between MethylCIBERSORT cell-type fractions and C-ECM scores. **E.** Heatmap of C-ECM genes in single-cell head-and-neck cancer RNAseq data. **F.** CAFs show the highest expression of C-ECM genes relative to other cell types in single-cell HNSCC data.

Finally, as an ultimate test of a CAF origin, we examined a dataset of single cell transcriptomes from head and neck cancers (GSE103322) ^13^ and found markedly higher expression of C-ECM genes in CAFs, which clustered together when the signature was projected onto the dataset (Figure 2E).

Indeed, C-ECM up and down ssGSEA scores were significantly elevated in CAFs compared to other cell types (Figure 2F), which we also independently verified in an additional colorectal cancer single-cell RNAseq dataset (GSE81861, Figure S2D) ^14^. Therefore, C-ECM profiles appear to be generated through the modulation of transcriptional profiles in CAFs specifically in malignancy.

Then, given that C-ECM scores correlate with CD8 T-cells and cytolytic activity (CYT) (Figure 2D and Figure S2C), and the fact that C-ECM up-scores are adversely prognostic despite the positive prognostic impact of CYT ^15^, we postulated that the C-ECM up-score may be enriched in immunologically ‘hot’ tumours, and our subsequent analyses uncovered robust evidence for this association using multiple orthogonal approaches. Accordingly, the C-ECM-up score was positively correlated with mutational burden (Rho = 0.23, p < 2.2e-16) while the down-signature was negatively correlated (Rho = −0.21, p < 2.2e-16) (Figure 3A).

**Figure 3:**
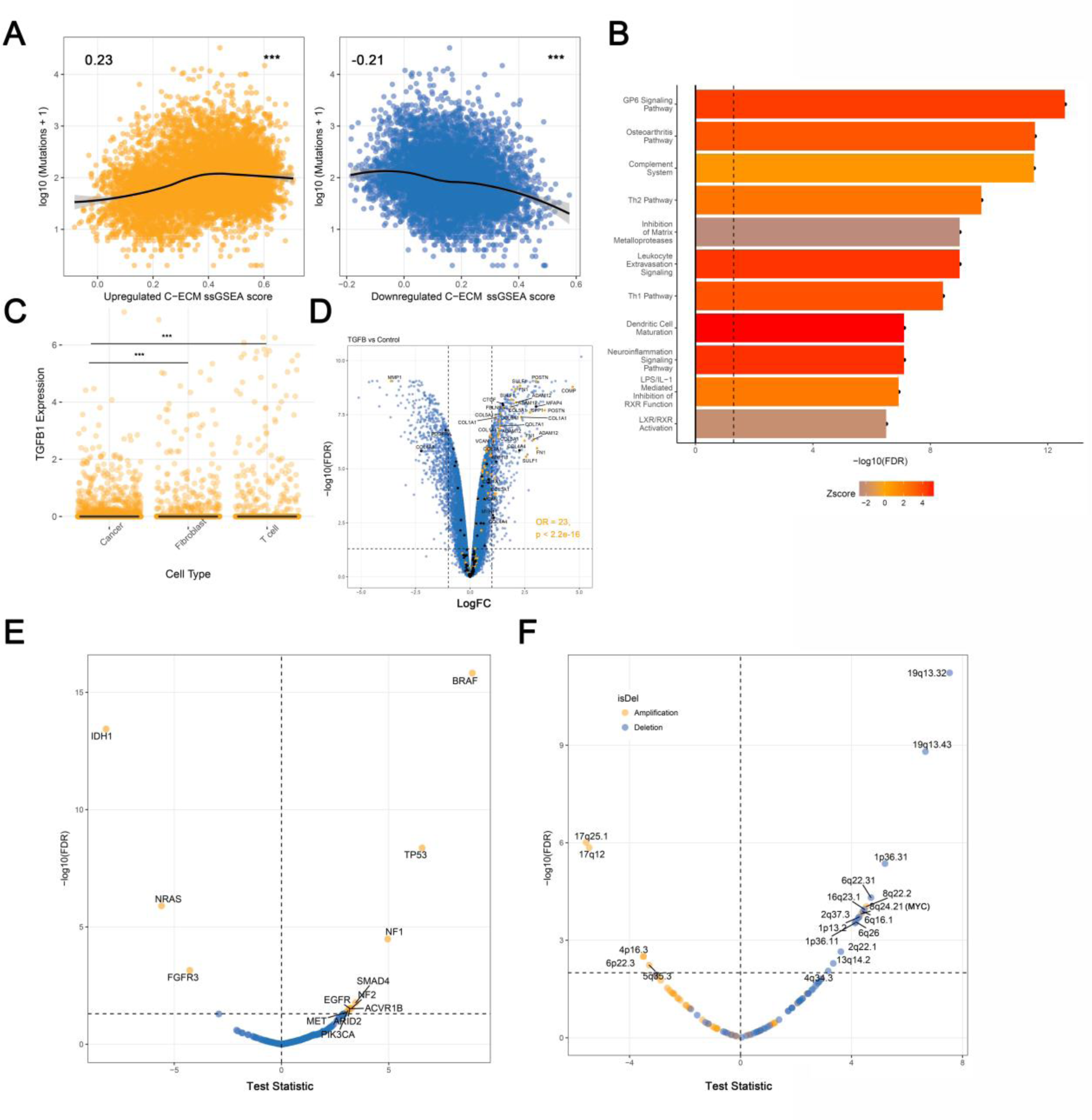
E-ECM scores are associated with immunologically hot tumours and TGF-β. **A.** ECM scores are significantly associated with mutational burden across cancer types. **B.** Canonical pathway analysis shows activation of inflammatory/adaptive-immune pathways. **C.** TGFB1 is significantly overexpressed in cancer cells in the single cell RNA-seq data. **D.** Volcano-plot showing enrichment for E-ECM genes in TGF-beta induced transcriptional changes in normal fibroblasts. **E and F** show linear model t-stats candidate mutational and copy-number alterations associated with ECM-up ssGSEA scores, adjusted for tumour type, on volcano-plots.

Associations between C-ECM scores and Class I neoantigen burden were also concordant (Rho = 0.21 and −0.21, p < 2.2e-16, Figure S3A) and so were associations between C-ECM scores and Microsatellite Instability, an immunotherapy biomarker *per se* ^16^ (Figure S3B). Additionally, we assessed macrophage polarisation using CIBERSORT ^17^ and found that the ECM-up signature was associated with a greater fraction of M1 relative to M2 (immunosuppressive) macrophages (Figure S3C). Finally, we found that multiple immune checkpoints, including *IDO1, B7-H3* and *PD-L2* were overexpressed in samples in the top quartile of the C-ECM up-score distribution relative to bottom quartile cancers after adjusting for tumour type (2FC, FDR < 0.01), indicating the upregulation of adaptive resistance mechanisms to immune-cell mediated destruction (Figure S3D). Moreover, these themes were broadly reinforced by IPA Canonical Pathway Analysis, which identified enrichment for inflammatory processes and adaptive immune responses enriched in samples in the top quartile of the C-ECM up-score (Figure 3B).

Next, since our data suggest that the C-ECM-up signature was generated by CAFs, and not by normal stroma, we endeavoured to find putative drivers responsible for this dysregulation. IPA Causal Network Analysis, after restriction to candidate regulators which by themselves differentially expressed between C-ECM-up top and bottom quartiles, identified TGF-β as one of the most activated regulators (Figure S3E) and upstream regulatory analysis further identified multiple *SMAD* transcription factors, *AP1* complex members that associate with SMADs ^18^, and *SMARCA4* ^19^ (Figure S3F), all critical for TGF-β transcriptional responses as activated in c-ECM-up-high cancers.

Moreover, orthogonal analyses using TCGA RPPA (Reverse Phase Protein Array) data (n = 4278), identified 13 differentially abundant peptides between upper and lower quartiles of the ECM-up score (FC > 1.3, FDR < 0.01, Figure S3G), most prominently, increased levels of Fibronectin and PAI1, both ECM components, with most showing associations with TGF-β (see Table S6), reinforcing the inference of activated TGF-β signalling. Indeed, in our RNA-seq analyses, TGF-β is significantly overexpressed in upper quartile C-ECM-up cancers along with multiple mediators of ECM deposition such as FGF family members (*FGF1*, *FGF18*), BMPs (*BMP1* and *BMP8A*) and the local sequestrators of TGF-β, *FBP1* and *LTBP1*. Moreover, in cancer cells in HNSCC single-cell RNAseq data (Figure 3C) it is overexpressed relative to fibroblasts and T-cells). Finally comparing the expression profiles of TGF-β treated immortalised ovarian fibroblasts (GSE40266) ^12^ versus untreated controls revealed marked enrichment for C-ECM genes amongst DEGs (Figure 3D), further buttressing the notion C-ECM gene dysregulation is a function of TGF- β signalling in CAFs.

As TGF-β is known to exert both pro-fibrotic and anti-proliferative effects, we decided to examine if enrichment for the C-ECM-up signature exerted specific adaptive constraints on the evolution of cancer genomes using TCGA data. Linear modelling implicated multiple genes after controlling for tumour type with known associations with TGF-β signalling from candidates positively selected in cancer ^20^.

Notable candidates included *TP53*, *SMAD4*, *BRAF*, *ACVR1B* and *NF1/2* (Figure 3E). We also implicated 18/111 significant GISTIC ^21^ peaks (Figure 3F), most notably *MYC* amplification (8q24.1) ( See Table S7 for detailed description of supporting literature), collectively confirming the hypothesized adaptation for TGF-β activation.

Finally, we tested whether C-ECM dysregulation is an immune evasion mechanism in the context of PD1/PD-L1 blockade, where immunologically ‘hot’ tumours are associated with responses ^22^. In two/three cohorts of PD-1 blockade ^23-25^, the C-ECM-up score was significantly higher in progressors (Figure 4A, p < 0.05, Wilcoxon’s Rank Sum Test). This was also true in pooled logistic regression accounting for cancer type, cytolytic activity, mutational load, a T-cell inflamed signature ^26^, cohort, antibody and prior anti-CTLA4 treatment (Figure 4B).

**Figure 4:**
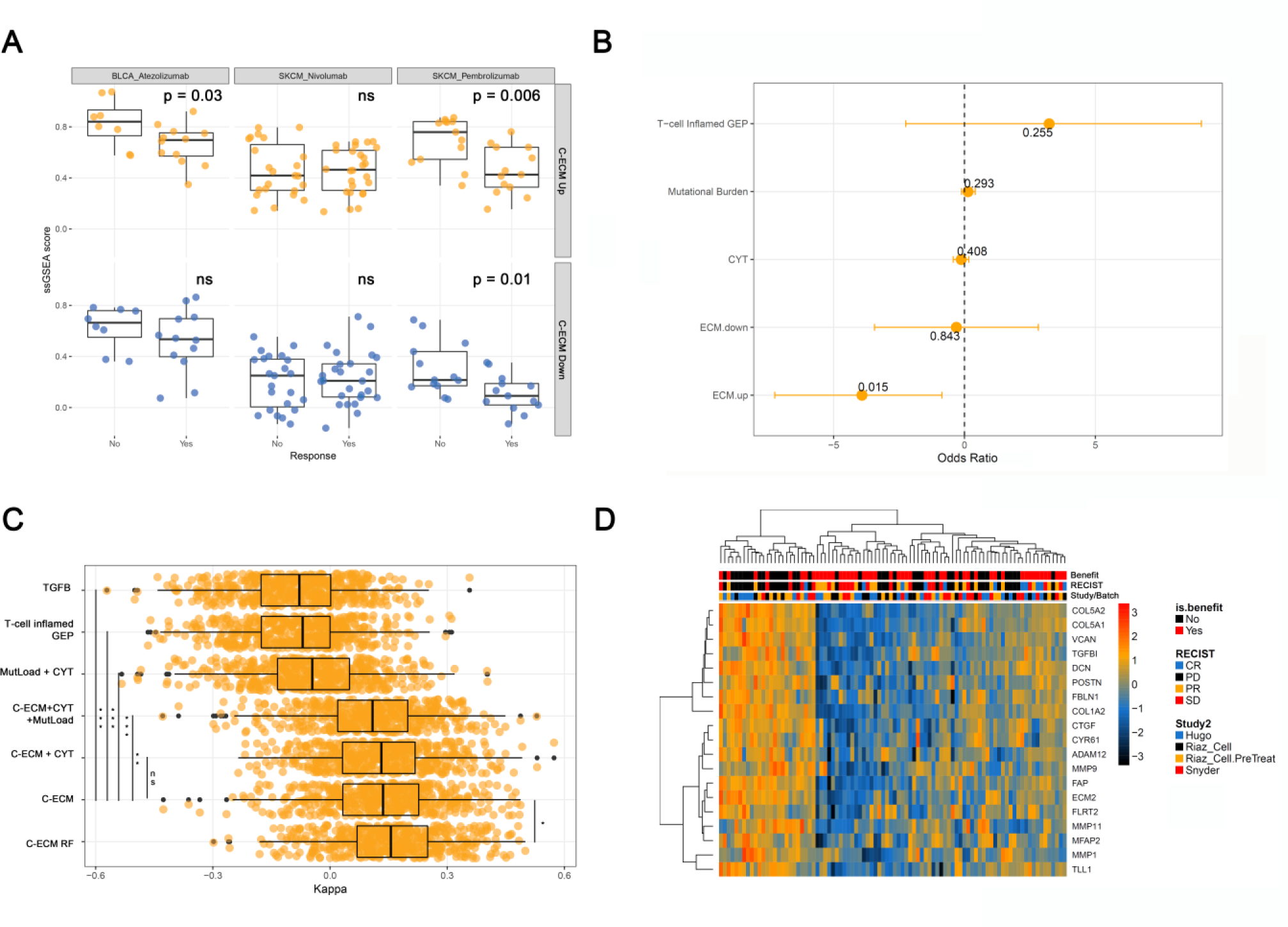
C-ECM scores predict failure of PD1-blockade. **A.** Boxplots showing distributions of C-ECM ssGSEA scores across multiple datasets of pretreatment biopsies from patients treated with PD1-blockade. Responders = CR/PR/SD. P.values from Wilcoxon’s Rank Sum Test. **B.** Coefficients from pooled logistic regression analysis evaluating various predictors on PD1-blockade response. **C.** Boxplots of Cohen’s Kappa from 0.632 bootstrapping (500 resamples), showing ECM-based models outperform other candidate biomarkers. Asterisks show q-values. **D.** Heatmap showing C-ECMGs differentially expressed between ICB responders and nonresponders after controlling for study-specific variation.

Next, comparing prediction performance using logistic regression with 0.632 bootstrapping ^27^ showed that models with C-ECM ssGSEA scores significantly outperformed those involving cytolytic activity, a T-cell inflamed signature, and mutation load alone (Figure 4C, S4A). Moreover, the aggregate score is comparable to a random forest fit with individual C-ECMs. Importantly, *TGFB1* expression alone does markedly worse than C-ECM based models, suggesting the presence of CAFs are required to convert *TGFB1* expression to an ICB-resistant phenotype through transcriptional modulation. Finally, restricted hypothesis testing using limma-trend found 19 C-ECM genes overexpressed at FDR < 0.1 (Figure 4D) between responders and nonresponders, defining a practical signature for clinical application (Figure S4B).

Given CAF-depletion *per se* is paradoxically associated with worse outcomes ^28^, approaches that seek to normalise the aberrant transcriptome in fibroblasts, possibly through TGF-β blockade, are likely to offer a promising route to boosting the efficacy of checkpoint blockade. Consistent with this, recent preclinical studies have uncovered evidence that simultaneous targeting of both TGF- β and PD-L1 can result in markedly better tumour control in multiple mouse models ^29^.

To summarise, we uncover a novel CAF-associated transcriptional pattern fundamentally linked to malignant transformation that permits immune evasion even in otherwise immunogenic tumours, explaining why signatures of negative selection in cancer may be so generally weak ^20^. In the process, we enhance our understanding of tumour-stromal interactions, and identify a key mediator of successful responses to PD1-blockade with significant translational implications.

**Figure legends – throughout, numbers on scatterplots indicate Spearman’s Rho, asterisks indicate statistical significance. * = p < 0.05, ** = p < 0.01, *** = p < 0.001. On all volcano plots, y axis = -log10 Fold Change, x axis = test statistic/ fold change/ Spearman’s Rho. On volcano plots, all enrichment statistics are from Fisher’s Exact Tests.**

